# Unveiling the Unique Structure and Singular Function of the Histone Deacetylase 2 (TgHDAC2) of *Toxoplasma gondii*

**DOI:** 10.1101/2025.03.21.644624

**Authors:** Caroline de Moraes de Siqueira, Julia Zamith Schwartz, Mariana Galvão Ferrarini, Mariana Sayuri Ishikawa Fragoso, Lysangela Ronalte Alves, Andrea Rodrigues Ávila, Tatiana de Arruda Campos Brasil de Souza, Sheila Cristina Nardelli

## Abstract

Histone deacetylases (HDACs) are enzymes traditionally recognized for their role in removing acetyl groups from lysines on histones. However, recent findings have revealed that many HDACs also target non-histone proteins. In *Toxoplasma gondii*, although we identified TgHDAC2, an enzyme annotated as a class I HDAC, we found that its substrates are non-histone proteins. Notably, TgHDAC2 possesses two unique peptide insertions within its HDAC domain, whose structural and functional roles were previously unknown. Using cross-linking mass spectrometry (XLMS), we resolved the three-dimensional structure of TgHDAC2, while biophysical analyses demonstrated that these insertions do not compromise the protein’s stability but play an important part in its function. Localization studies revealed differential expression of TgHDAC2 throughout the cell cycle, with prominent enrichment around daughter cells during mitosis and cytokinesis. Its deletion severely disrupts parasite replication, suggesting a critical role in cell cycle regulation. RNA sequencing of TgHDAC2 knockout parasites highlighted significant downregulation of genes involved in membrane composition, cytoskeletal organization, and cell signaling pathways, further supporting its role in modifying non-histone proteins. Collectively, our results suggest that TgHDAC2 acts as a deacetylase for non-histone proteins, modulating cytoskeletal and membrane proteins critical for *T. gondii* cell cycle progression and replication.

**IMPORTANCE:** *Toxoplasma gondii* is an obligate intracellular parasite and a significant global public health concern. It is estimated that up to one-third of the world’s population may be infected, depending on the region, with even higher prevalence rates in South America due to the circulation of atypical and more virulent strains. Understanding the biology of this parasite and identifying novel therapeutic targets is therefore critical, as current treatments are outdated and ineffective against the chronic phase of toxoplasmosis. In this study, we identified a novel lysine deacetylase that plays an essential role in *T. gondii* replication, highlighting its potential as a promising therapeutic target.

## 1 Introduction

Proteins are subject to a diverse array of biochemical modifications, including phosphorylation, acetylation, methylation, glycosylation, and others. These alterations, collectively referred to as post-translational modifications (PTMs), are essential for modulating protein structure and function. PTMs influence gene expression, direct the transport of proteins to specific cellular compartments, regulate protein-protein interactions, and play pivotal roles in cell signaling pathways (1).

Lysine acetylation is a prevalent and reversible modification regulated by histone acetyltransferases (HATs) and histone deacetylases (HDACs). These enzymes respectively catalyze the addition or removal of acetyl groups on both histone and non-histone targets (2).

The modification of lysines through acetylation is well known for its role in regulating gene transcription via histones, the primary substrates of HDACs (3). Beyond histones, acetylation of non-histone proteins highlights the diversity of this modification, which is involved in processes such as metabolism, cytoskeleton organization, protein folding, centrosome regulation, translation, and RNA processing (4) (5) (6) (7). The mechanisms by which HATs and HDACs select their non-histone substrates need to be better understood (2). HATs and HDACs are often complexed with other proteins and may associate with multiple complexes with distinct functions, site specificities, and enzyme activities (8). The molecular effects of acetylation residues within non-histone proteins are still emerging (9) (10).

HDACs are classified into Zn^2+-^dependent histone deacetylases (classical HDACs) and NAD^+^-dependent sirtuins (2).

Historically, HDACs have been key targets for the development of inhibitors aimed at treating cancer and other diseases, primarily due to their role in regulating gene expression. A broad range of small-molecule inhibitors targeting HDACs have been identified as potential starting points for new treatments for parasitic diseases (11). Inhibitors of HDACs have been studied in protozoan parasites, such as *Plasmodium* and *Leishmania* (12) (13). In *Toxoplasma gondii*, two compounds targeting HDACs have been identified. The compound FR235222 effectively inhibits *T. gondii* growth by specifically blocking TgHDAC3 (14). More recently, another compound, MC1742, was discovered as a potent HDAC inhibitor (HDACi) targeting histone HDACs, significantly affecting parasite growth (15).

*Toxoplasma gondii* is an apicomplexan parasite and the causative agent of toxoplasmosis. This cosmopolite disease is hazardous in immunocompromised patients and pregnant women. According to the ToxoDB database (http://ToxoDB.org), this species has seven identified HDACs (five classical HDACs and two sirtuins), most of which are essential for parasite survival, as determined through genetic screening using CRISPR-Cas9 (16). Among these, only TgHDAC3 and TgKDAC4 have been described. TgHDAC3 is part of a co-repressor complex involved in deacetylating histone H4 at transcription initiation sites during the differentiation of tachyzoites into bradyzoites, thereby activating the expression of bradyzoite-specific genes (14). TgKDAC4, located in the apicoplast, functions in protein deacetylation within this organelle, independent of histones (17).

Apart from TgHDAC3 and TgKDAC4, the function of other HDACs in *Toxoplasma* remains to be determined. This includes TgHDAC2, a class I HDAC whose HDAC domain is unique to species within the family Sarcocystidae. Notably, TgHDAC2 contains two large peptide insertions within its HDAC domain (18). These insertions, equivalent to 20 kDa are found only in a few apicomplexan species, including *Toxoplasma*, *Neospora*, and *Hammondia*, and their functions and structure remain unknown. Overall, the HDACs of this parasite exhibit regions with extensions and insertions that may contribute to a distinct mode of action (19).

In this study, we characterized the function of TgHDAC2 through a classical knockout approach, revealing significant defects in replication. These defects impaired S-phase progression, leading to disorganization within the parasitophorous vacuole. We also performed a structural analysis that suggests that the unique insertions in TgHDAC2 contribute to its function rather than structure stability. Finally, we proposed a potential structure for TgHDAC2 using cross-linking mass spectrometry (XLMS). Collectively, our data indicate that this protein targets non-histone substrates, including components of the membrane and cytoskeleton.

## 2 Material and Methods

### 2.1 Parasite culture

*Toxoplasma gondii* RHΔhxgprtΔku80 strain was grown in normal human dermal fibroblasts (NHDF; Lonza, Basel, Switzerland) using Dulbecco’s Modified Eagle Medium (DMEM) supplemented with 10% fetal bovine serum, 2 mM L- glutamine, and 100 μg/mL of penicillin/streptomycin. Cultures were maintained at 37°C in an atmosphere containing 5% CO2. Parasites were harvested and lysed using 27G needles for syringe lysis, followed by washing with cytomix buffer (2 mM EDTA, 120 mM KCl, 0.15 mM CaCl2, 10 mM K2HPO4/KH2PO4, 25 mM HEPES, and 5 mM MgCl2). After washing, parasites were electroporated with a Nucleofector II electroporator (Amaxa Lonza, Basel, Switzerland) using the T16 program with a single pulse.

### 2.2 Generation of knockout and complemented parasites

A 1kb sequence corresponding to the downstream and upstream untranslated regions (UTRs) from the *tghdac2* (TGME49_249620) was amplified using genomic DNA from the *T. gondii* RHΔhxgprtΔku80 strain and conjugated to the hypoxanthine-xanthine-guanine-phosphoribosyl transferase (*hxgprt*) gene via fusion PCR (20). The resulting cassette was transfected into *T. gondii* RHΔhxgprtΔku80, and the knockout parasites were selected by adding 50 μg·ml-^-1^ xanthine and 25 μg·ml^-1^ mycophenolic acid to the culture. Parasite clones were then isolated through limiting dilution (21).

The sequence corresponding to the *tghdac2* gene was amplified to generate complemented parasites using cDNA from the *T. gondii* RH*ΔhxgprtΔku80* strain. Primers were designed to include 40 base pairs of homologies to the uracil phosphoribosyl transferase (*UPRT*) gene. The *tghdac2* gene was inserted into the genome by transfecting the amplicon along with a CRISPRCas9 plasmid containing a single guide RNA (sgRNA) targeting the UPRT locus (22). Resistant parasites were selected by adding 10 μM of floxuridine (FUDR) to the culture. Resistant clones were subsequently isolated through limiting dilution, and PCR confirmed the successful reintroduction of *tghdac2*.

### 2.3 Plaque assay

A total of 500 parasites from -control (RHΔ*hxgpr*tΔ*ku80*), knockout (RHΔ*tghdac2*), and TgHDAC2-complemented (RHΔ*tghdac2*::*tghdac2*) strains were used to infect confluent NHDF cells cultured in 6-well plates. After five days of incubation, the cells were fixed with ice-cold 100% methanol and stained with 50μL/mL Giemsa solution (Sigma-Aldrich, San Luis, Missouri, USA) for 20 minutes to visualize lysis plaques (23). The assay was performed in three independent experiments, and statistical analysis was conducted using one-way analysis of variance (ANOVA) followed by Tukey’s test.

### 2.4 Replication assay

The equivalent of 1 x 10^5^ tachyzoites from RH*ΔhxgprtΔku80*, RHΔ*tghdac2*, and RHΔ*tghdac2*::*tghdac2* strains were used to infect NHDF cells cultured in 24-well plates, each containing one coverslip per well. After 24 hours of incubation, the infected cells were washed with 1x PBS and fixed with 4% paraformaldehyde. The parasites were visualized by indirect immunofluorescence using rabbit anti-IMC1 antibodies (1:1,000 dilution, kindly provided by Gary Ward, University of Vermont).

### 2.5 Cell cycle analysis

Intracellular RH*ΔhxgprtΔku80*, RHΔ*tghdac2*, and RHΔ*tghdac2*::*tghdac2* parasites were collected from infected cells after 24 and 48 hours of culture via manual lysis using repeated passages through a 27-gauge needle. The cell lysates were washed twice with PBS, and parasites were filtered using a 3μm Whatman^®^ membrane to minimize cell debris. The parasites were fixed in 70% ethanol and stored at -20°C for at least 18 hours.

Following fixation, the samples were centrifuged at 3,000 x g for five minutes at 4°C to remove ethanol. Parasites were resuspended in 200 μL of staining solution containing 10 μg/mL propidium iodide (PI), 100 μg/mL RNase A, and 0.1% Triton X-100 and incubated for 30 minutes at room temperature in the dark. PI-stained nuclei were analyzed by flow cytometry using a FACSAria^TM^ flow cytometer (BD Biosciences, Franklin Lakes, New Jersey, USA). Gates were defined to exclude debris and cell clumps, and the cell cycle profiles were analyzed using FlowJo software™ v10.8 Software (24) (BD Life Sciences, Franklin Lakes, New Jersey, USA).

### 2.6 Anti-TgHDAC2 production

Polyclonal antibodies against TgHDAC2 were generated using the peptide sequence (GRRKSVLYFYDENIC) as the antigen, synthesized by FastBio (Ribeirão Preto, São Paulo, BR). Two rabbits were immunized, and antibody production was performed using the PolyExpress^TM^ method (GenScript, Piscataway, New Jersey, USA).

### 2.7 Western blot

Freshly lysed parasites (RH*ΔhxgprtΔku80*, RHΔ*tghdac2*, and RHΔ*tghdac2*::*tghdac2*) were prosseced for Western Blot following standard procedures (25). The primary anti-TgHDAC2 antibody (1:50) was incubated overnight at 4°C. The membrane was then incubated for one hour with a secondary anti-rabbit antibody (1:10,000), conjugated to peroxidase. The signal was developed using chemiluminescence (SuperSignal West Pico PLUS, Thermo Fisher Scientific, Waltham, Massachusetts, USA) and visualized with the iBright Imaging System (Thermo Fisher Scientific, Waltham, Massachusetts, USA).

### 2.8 Immunofluorescence microscopy

NHDF cells grown on coverslips were infected with *T.gondii* as described above, then fixed with 4% paraformaldehyde for 20 minutes. The cells were subsequently permeabilized with 0.25% Triton X-100 and blocked with 1% PBS/ BSA. After blocking, the cells were incubated with the primary antibodies mouse anti-IMC1 (1:1,000; kindly provided by Dr. Peter Bradley) or rabbit anti-IMC1 (1:1,000) for one hour or anti-TgHDAC2 (1:25) overnight, followed by incubation with Alexa secondary antibody (Invitrogen, Waltham, Massachusetts, USA) 488 or 546 at a 1:600 dilution. Along with the secondary antibody, the coverslips were incubated with 10 μM DAPI (4’,6-diamidino-2 phenylindole). The slides were mounted with N-propyl galactoside to preserve fluorescence and analyzed using a Leica DMI6000 B fluorescence microscope (Leica, Wetzlar, Germany).

### 2.9 Scanning electron microscopy

NHDF cells, previously grown on coverslips, were infected with RH*ΔhxgprtΔku80*, RH*Δtghdac2*, and RH*Δtghdac2::hdac2* strains for 24 hours. After this period, the cells were fixed for 30 minutes with 2.5% glutaraldehyde and 4% paraformaldehyde (PFA) diluted in 0.1 M cacodylate buffer (pH 7.2), followed by two washes for five to 10 minutes with the same buffer. The cells were then incubated for one hour with 1% OsO4 diluted in 0.1 M cacodylate buffer. After incubation, the samples were washed three times with the same buffer and then dehydrated in increasing acetone concentrations for 10 minutes. Critical point drying was performed, followed by gold sputter-coating. The cell surface was then detached using an adhesive tape.

### 2.10 Transcriptome of mutant parasites

The RH*ΔhxgprtΔku80*, RH*Δtghdac2*, and RH*Δtghdac2::hdac2* parasites were collected from infected host cells by lysing through passages using a 25-gauge needle. The cell lysates were washed twice with PBS, and the parasites filtered through a 3μm Whatman^®^ membrane (Maidstone, United Kingdom) to reduce cell debris. Total RNA was isolated using the RNeasy kit (Qiagen, Redwood City, California, USA) according to the manufacturer’s protocol. Following the manufacturer’s instructions, 1 μg of RNA was used for mRNA library preparation with the Illumina Stranded mRNA Prep Ligation kit.

Sequencing reads were mapped against the *T. gondii* ME49 genome (NC_031467.1) using STAR v2.7.10a (26) with default parameters. Gene counts for uniquely mapped reads were obtained with featureCounts v2.0.1 from the Subread package (27). Differentially expressed genes (DEGs) were identified using the DESeq2 v1.34.0 package (28) within R (29), with an adjusted p-value of less than 0.05. Functional characterization of DEGs was performed using Omics Box software (30) and functional enrichment analysis was conducted with clusterProfiler within R (31). Data were plotted using the ggplot2 package in R (32). The raw RNA-seq data have been deposited in the NCBI Sequence Read Archive (SRA) under BioProject ID PRJNA1233816 and SRA temporary accession number SUB15162896.

### 2.11 Plasmid construction

The TgHDAC2 and TgHDAC2^Δ74-276^ (ToxoDb gene ID TGME49_249620) genes were amplified by PCR using complementary *T. gondii* DNA. For the full-length TgHDAC2 gene, attB sites were incorporated into specific primers to amplify the entire coding region. The amplicon was then cloned into the pDONR221 vector (Invitrogen, Waltham, Massachusetts, USA) via recombination, following the manufacturer’s instructions. It was subsequently subcloned into the pDEST17 expression vector (Invitrogen, Waltham, Massachusetts, USA).

The cDNA of *T. gondii* was also used as a template to clone TgHDAC2^Δ74-276^. In this case, two amplifications were performed: one upstream of the insert, where the restriction sites for *Nde*I and *Bam*HI were added, and the other downstream of the insert, where the restriction sites for *Bam*HI and *Hind*III were incorporated. Both regions were then cloned into the pET28a expression vector (Novagen, Burlington, Massachusetts, USA).

### 2.12 Protein expression and purification

TgHDAC2 protein was expressed in transformed *E. coli* Rosetta gami^TM^ 2 (DE3) pLysS by induction with 0.5 mM IPTG after reaching an OD600 of 0.4, followed by incubation for four hours at 37°C under shaking conditions. For TgHDAC2^Δ74-276^, the *E. coli* ΔSlyD strain was used. When the OD600 reached 0.4, 0.5 mM IPTG was added to induce protein expression for 16 hours at 20°C under agitation.

After induction, the samples were harvested by centrifugation at 10,000 x g for 10 minutes at 4°C, and the supernatants were discarded. The pellet was resuspended in 50 µl of sonication buffer (20 mM sodium phosphate, pH 7.5, and 500 mM NaCl), containing the protease inhibitors 0.1 M phenylmethylsulfonyl fluoride (PMSF) and 5 mM benzamidine, and 10 µg/ml lysozyme. The cell suspension was subjected to lysis by sonication, with eight cycles lasting 15 seconds each at an amplitude of 5. The protein extract was then sedimented by centrifugation at 25,000 x g for 30 minutes, and the pellet was resuspended in 200 mM sodium phosphate buffer A (pH 7.5), supplemented with 8 M urea and 500 mM NaCl.

Purification was performed using affinity chromatography on an AKTA pure chromatography (GE Healthcare, Chicago, Illinois, USA), with a column containing 1 mL HisTrap HP (GE Healthcare, Chicago, Illinois, USA) nickel resin. Elution was conducted with a linear gradient of buffer B (20 mM sodium phosphate, pH 7.5, 500 mM NaCl, 1 M imidazole, and 4 M urea). Eluted proteins were refolded via dialysis under gradually decreasing urea concentrations until the urea was completely removed. A second round of affinity chromatography was performed under the same conditions as the initial purification.

### 2.13 Circular dichroism (CD)

For CD and thermal denaturation experiments, the purified protein concentrations were set at 14.5 µM (TgHDAC2^Δ74-276^) and 2.49 µM (TgHDAC2), followed by dialysis in a 20 mM sodium phosphate buffer (pH 7.5) containing 180 mM NaCl. The samples were placed in a 1 mm optical path cuvette in a Jasco J815 CD spectrometer (Jasco Corporation, Hachioji, Tokyo, Japan), equipped with Spectra Measure for spectral acquisition. Wavelengths from 198 to 260 nm were measured. For thermal denaturation, the temperature was gradually increased from 20°C to 90°C, and the CD signal at wavelengths 205, 208, 215, and 222 nm was recorded after each 5°C increment. Upon reaching 90°C, a continuous spectrum was collected between 190 and 260 nm.

### 2.14 Cross-linking mass spectrometry (XLMS)

The crosslinking experiment with purified TgHDAC2 was performed as previously described (32). Briefly, 10 mg/ml of DSS crosslinker was added to 200 µg of purified protein and incubated at 27°C for two hours. The reaction was then stopped by adding 100 mM ammonium bicarbonate (pH 8) and 10 mM DTT. The sample was heated at 60°C for 30 minutes and then cooled to room temperature. Iodocetamide was added to the sample to a final concentration of 30 mM, and the mixture was incubated in the dark for 30 minutes. The sample was digested overnight with trypsin (1:20 w/w) at 37°C. After digestion, trifluoroacetic acid (TFA) was added for sample acidification and re-quantification. A total of 10 µg of digested protein were used for each stage tip in the C18 desalting step. The column was activated with 100 µL of methanol, centrifuged at 1,000 x g for two minutes, and then equilibrated twice with 0.1% formic acid. The sample was added to the column, centrifuged at 1,000 x g for five minutes, and this process was repeated until the entire sample had passed through the column. The stage tip was then washed twice with 0.1% formic acid. Analysis was performed using a Lummus mass spectrometer and SIMXL software to generate de cross-linkers map (33). The map obtained from SIMXL was submitted to iTasser (34) to build the model.

### 2.15 *In silico* Analysis

For the *in silico* analysis of potential post-translational modifications (PTMs), the ScanProsite server (Expasy) (35) was used to identify conserved motifs within TgHDAC2. Additionally, functional predictions were performed using the Profunc analysis tool (36), which assigned potential functions to TgHDAC2 based on structural similarities.

## 3 Results

### 3.1 *Toxoplasma gondii* has sequences inside the HDAC domain that are not involved in the protein’s stability

TgHDAC2 (http://TGME49_249620-ToxoDB.org) exhibits unique characteristics. The protein has a molecular weight of approximately 67 kDa, with an HDAC domain encompassing nearly the entire protein. Within the HDAC domain, two peptide insertions are present: the first consists of 73 amino acids, followed by 12 conserved amino acids characteristic of the HDAC2 domain, and the second insertion comprises 117 amino acids (Fig 1B). The function of these insertions is currently unknown.

**Figure 1.**
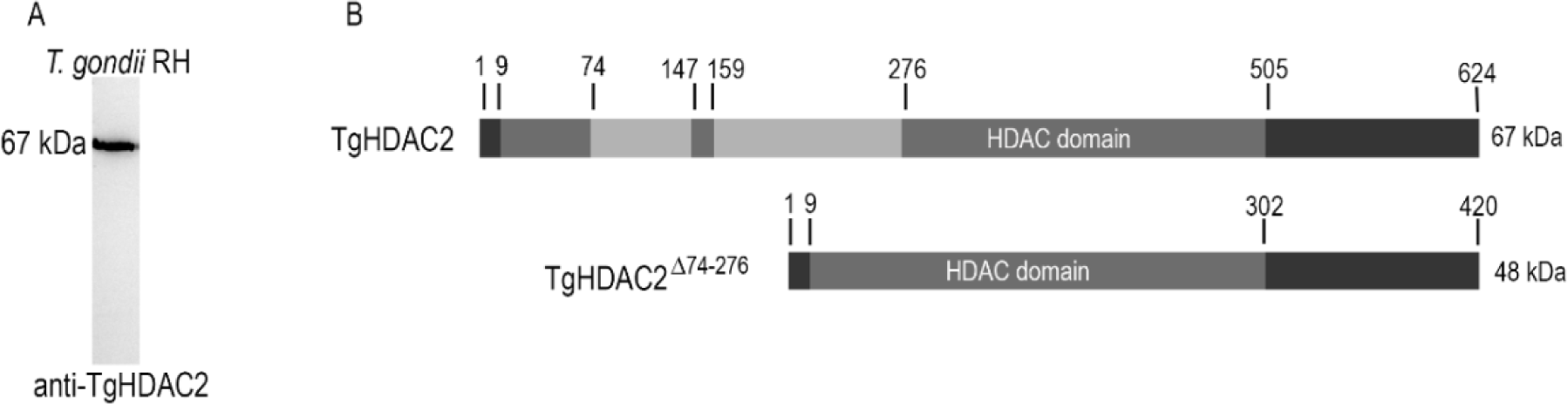
(a) Western blot of *T. gondii* extract probed with a specific anti-TgHDAC2 polyclonal antibody. The full-length TgHDAC2 protein is expressed in tachyzoite forms of this parasite, with Western blot analysis revealing a band at the expected size of 67 kDa. (b) The full-length protein consists of 624 amino acids (dark gray). The HDAC domain (gray) spans amino acids 9 to 505, interrupted by two peptide insertions (light gray) between amino acids 74 and 147 and between amino acids 156 and 276. The construct lacking the amino acid insertions (TgHDAC2 ^Δ74-276^) results in a protein of 420 amino acids (dark gray) and a molecular weight of 48 kDa, with a complete HDAC domain (gray) between amino acids 9 and 302.

Our initial hypothesis was that the peptide insertions could serve as a regulatory domain where PTMs might be introduced to modify the protein’s activity. To investigate potential PTMs, *in silico* analysis using the ScanProsite server (Expasy) identified several phosphorylation sites of protein kinase C (PKC) and casein kinase II (CK2) within the peptide insertions and the C-terminal portion (Table 1). The Expasy analysis also predicted sites for amidation, phosphorylation y cAMP- and cGMP-dependent protein kinases, N-myristoylation, and N-glycosylation. These findings suggest that the PTMs in these regions play a role in modulating the stability or activity of TgHDAC2.

**Table 1.** PTM sites predicted for TgHDAC2 in full form. Several potential PTM sites were identified (Table 1), with some located within the peptide insertions. This suggests that peptide insertions contribute to the regulation of enzyme activity through PTMs. (*) Confirmed sites.

| PTM type | Site | PTM found inside insertions? |
| --- | --- | --- |
| **Amidation** | 1–4 | No |
|  | 329–332 | No |
| **cAMP- and cGMP-dependent protein kinase phosphorylation site** | 3–6 | No |
| **Casein kinase II phosphorylation site** | 78 | Yes |
|  | 154 | No |
|  | 171 | Yes |
|  | 207 | Yes |
|  | 222 | Yes |
|  | 274 | Yes |
|  | 537 | No |
|  | 558 | No |
|  | 579 | No |
|  | *602 | No |
| **Protein kinase C phosphorylation site** | *85 | Yes |
|  | 198 | Yes |
|  | 244 | Yes |
|  | 378 | No |
|  | 439 | No |
|  | 492 | No |
|  | 558 | No |
|  | 619 | No |
| **N-myristoylation site** | 231–236 | Yes |
|  | 240–245 | Yes |
|  | 253–258 | Yes |
|  | 304–309 | No |
|  | 326–331 | No |
|  | 370–375 | No |
|  | 403–408 | No |
|  | 407–412 | No |
|  | 419–424 | No |
|  | 488–493 | No |
|  | 567–572 | No |
| **N-glycosylation site** | 463 | No |

While certain PTMs have been predicted, one has been confirmed at position 85 (within the amino acid insertion). These amino acid insertions may also play a crucial role in enhancing the overall protein’s stability. To evaluate the contribution of these insertions to TgHDAC2 stability, we obtained the recombinant protein derived from the full-length TgHDAC2 sequence, as well as its counterpart lacking the amino acid region between residues 74 and 276 (TgHDAC2^Δ74-276^), as shown in Fig. 1.

CD was used to assess the biophysical properties of both protein forms, providing insights into their secondary structure. The CD spectra for TgHDAC2 and TgHDAC2^Δ74-276^ demonstrated that both proteins are structured, exhibiting characteristics of proteins composed of α-helices and β-sheets (Fig. 2). While TgHDAC2 displayed a negative signal at 198 nm, the signal was not as pronounced as it would be expected for a random coil.

**Figure 2.**
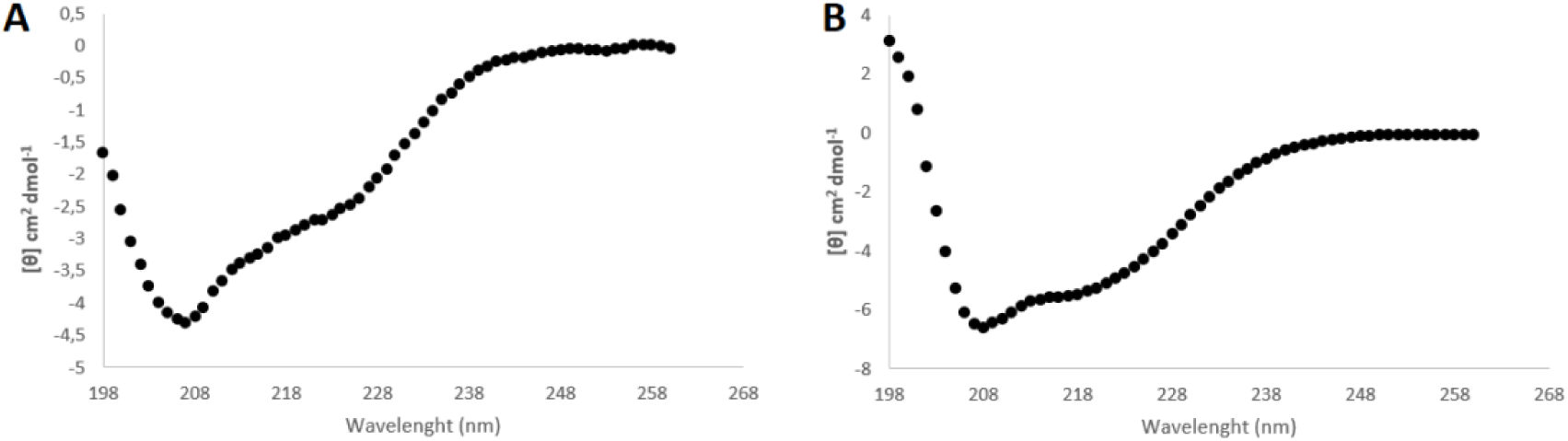
Circular dichroism spectra for TgHDAC2 (A) and TgHDAC2 ^Δ74-276^ (B) recorded at 20°C over a wavelength range of 198–258 nm. It is possible to note that both proteins are structured, with negative signals at 222 nm and 208 nm and a positive trend at 198nm.

After confirming that both proteins are structured, their stability was assessed using thermal denaturation. This method assesses the ability of proteins to renature after exposure to temperatures up to 90°C, providing a reliable measure of proteinprotein stability. The analysis revealed that TgHDAC2^Δ74-276^ is more stable than full-length TgHDAC2 (Fig. 3). Unlike TgHDAC2^Δ74-276^, the full-length TgHDAC2 is unable to renature and return to its initial conformation following denaturation. These findings suggest that peptide insertions do not contribute to the structural stability of this protein. However, while this region does not contribute to the stability of TgHDAC2, it is possible that the insertions are functionally significant rather than structurally essential. To further investigate how the insertions might influence the protein’s structure and potentially its function, we proceeded to analyze the tertiary structure of TgHDAC2.

**Figure 3.**
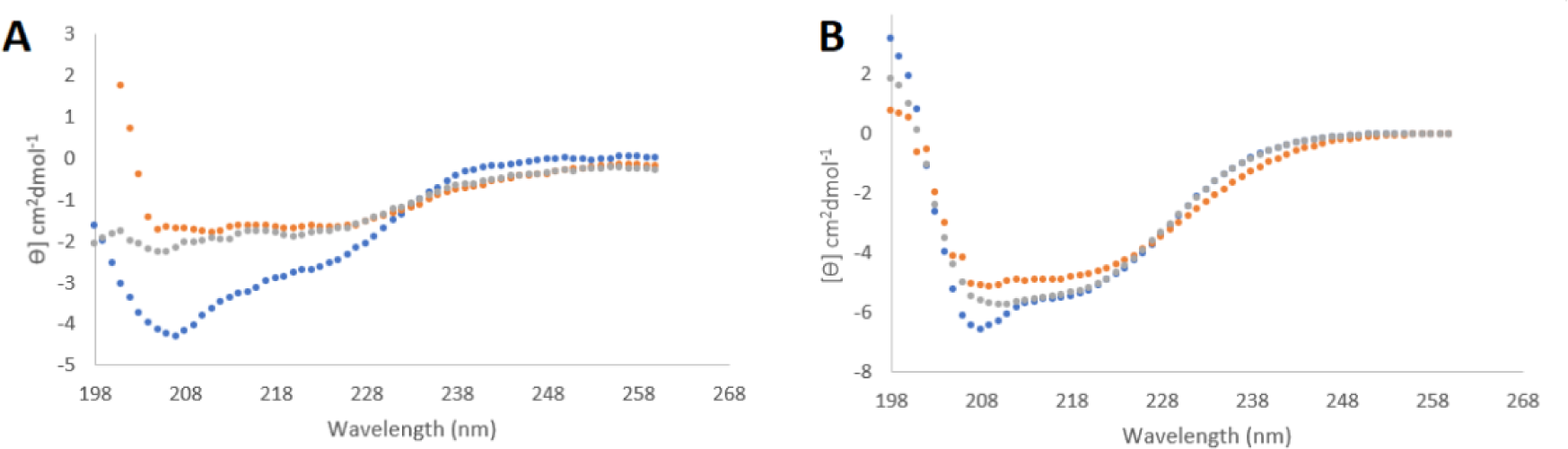
Circular dichroism spectra of TgHDAC2 (A) and TgHDAC2^Δ74-276^ (B) recorded at different temperatures: 20°C (blue), 90°C (orange), and 20°C after having heating to 90°C (grey), over a wavelength range of 198–268 nm. It is possible to observe that TgHDAC2 shows a significant loss of structure after heating, with limited renaturation. In contrast, TgHDAC2^Δ74-276^ exhibits less pronounced structural loss and efficient renaturation.

### 3.2 TgHDAC2 peptide insertions are predominantly composed of α-helix

To investigate the three-dimensional structure of TgHDAC2, XLMS was performed using DSS to crosslink the purified full-length protein, allowing identification of residues in close spatial proximity. A total of 63 crosslinked peptides were identified, and a map of interactions was made. Based on the spatial restraints obtained from X-linkers, five structural models were generated. These models exhibited differences of up to 1Å in RMSD, indicating convergence toward a similar structure. Among these, model 5 was identified as the most consistent with the majority of experimental DSS XLinkers, as summarized in Supplemental Table 1.

The peptide insertion comprises α-helices that intricately interact with the HDAC domain (Fig. 4). Notably, the α-helix formed by residues 215–219 appears to close the funnel of the HDAC domain, serving as a regulatory protein element (Fig. 4). Detailed interactions between the peptide insertion and the HDAC domain are provided in Supplemental Table 1. The interface between the two structures is stabilized by 41 hydrogen bonds and 24 salt bridges, yielding a stable interface with a predicted delta G of -20.0 kcal/mol. These findings underscore the robustness of the interaction.

**Figure 4.**
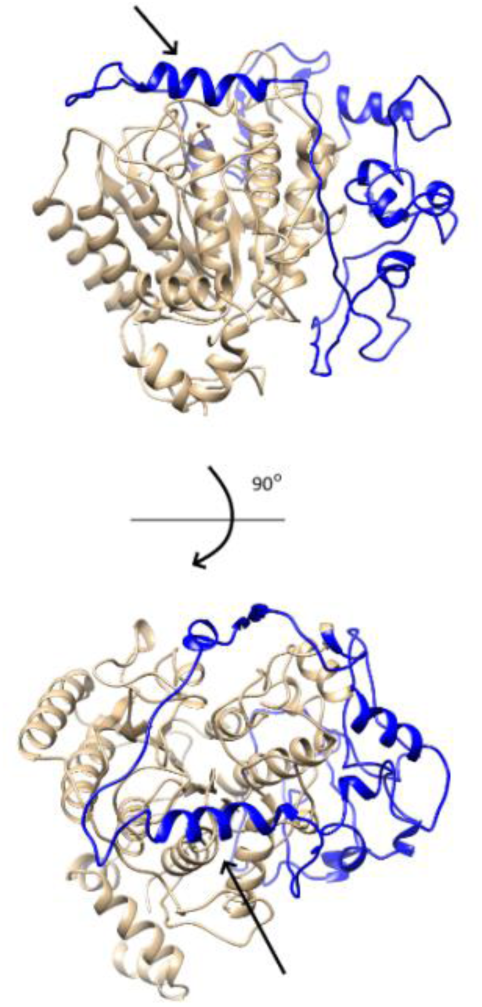
The three-dimensional structure of TgHDAC2 was determined through XLMS and molecular modeling. The peptide insertion is highlighted in blue, while the HDAC domain is depicted in wheat. The arrow indicates the helix encompassing residues 215-219.

Subsequently, this model was applied in Profunc analysis, which identified potential functions for TgHDAC2 with e-values ≤ 0.083. The predicted roles include cyclin-G-associated kinase, microtubule crosslinking factor 1, serine/arginine repetitive matrix protein 2, myb-related transcription factor, and profilin partner.

Having established its distinct structural features, the next step is to explore the functional implications of these findings. In this regard, we proceeded with the functional characterization of TgHDAC2 to gain a deeper understanding of how these structural attributes contribute to its role within the parasite.

### 3.3 TgHDAC2 is located at the membrane of *Toxoplasma* and is essential for replication and virulence

To identify the localization of TgHDAC2, we ordered a polyclonal antibody (FastBio). While class I HDACs in metazoans are typically nuclear proteins, TgHDAC2 is an exception. The polyclonal antibody was used in immunofluorescence experiments, detecting the protein outside the nuclei. TgHDCA2 is localized in dots prior to division and at the membrane during mitosis and cytokinesis (Fig. 5A), with minimal detection observed during interphase. Colocalization with anti-inner membrane complex (IMC; provided by Peter Bradley) confirmed its localization at the membrane. This antibody, originally produced for *Neospora*, cross-reacts with IMC in *Toxoplasma*, though the specific IMC recognized remains unclear (37) (Fig. 5B).

**Figure 5.**
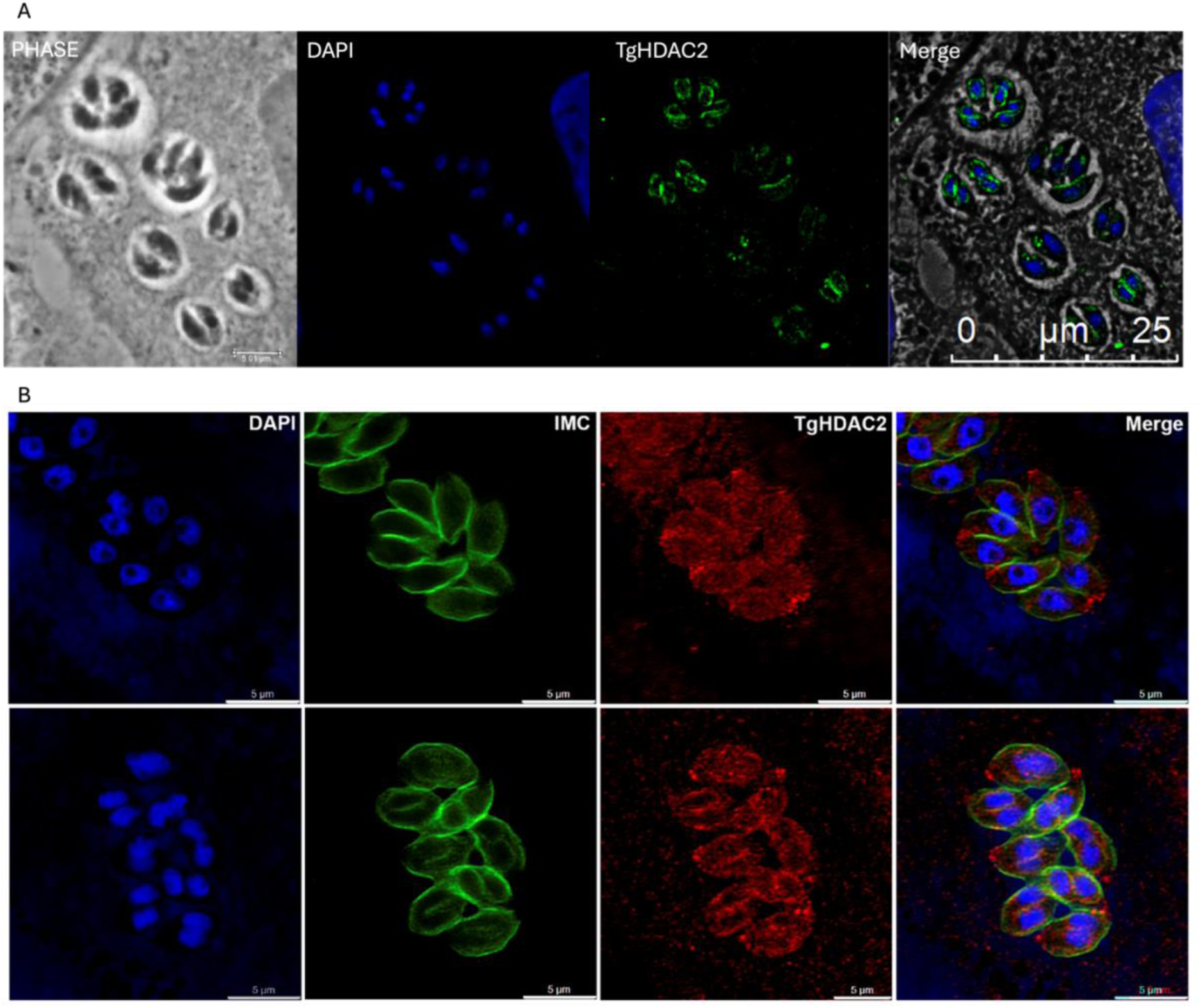
TgHDAC2 is expressed in replicating parasites. (A) HFF cells were infected with RH *ΔhxgprtΔku80* parasites, and IFA using an anti-TgHDAC2 polyclonal antibody revealed that TgHDAC2 is present during early parasite budding and is more prominent in replicating parasites. (B) Co-localization with anti-IMC antibodies confirms that TgHDAC2 is localized to the daughter cell IMC compartment. DAPI was utilized to visualize the nucleus.

The absence of nuclear localization for TgHDAC2 suggests its role in the acetylation of non-histone proteins. To investigate the function of TgHDAC2, we generated a classical knockout by using approximately 1,000 pb of the 5’ UTR and 3’ UTR of the gene, which were then fused to the selection marker *hxgprt*. The cassette was transfected into RH ΔhxgprtΔku80 parasites. After selection, clones were confirmed by PCR and Western blot analysis (Supp. Fig.1).

The effect of *tghdac2* absence is evident in plaque assay experiments, where the knockout parasites formed fewer plaques (Fig. 6A-B), indicating a defect in virulence or replication. To confirm these findings, the *tghdac2* gene was complemented using CRISPR-Cas9. The insertion was confirmed by PCR and Western blot analysis (Supp. Fig. 2). Immunofluorescence confirmed the replication defect, as knockout parasites displayed fewer parasites per vacuole on average (Fig. 6C-D). We then analyzed whether the replication defect was due to a delay in the cell cycle using flow cytometry (Fig. 7A-B). A higher number of knockout parasites (*Δtghdac2*) were observed in the S and M phases after 48 hours of culture compared to the wild type, confirming a replication-related defect (Fig. 7C).

**Figure 6.**
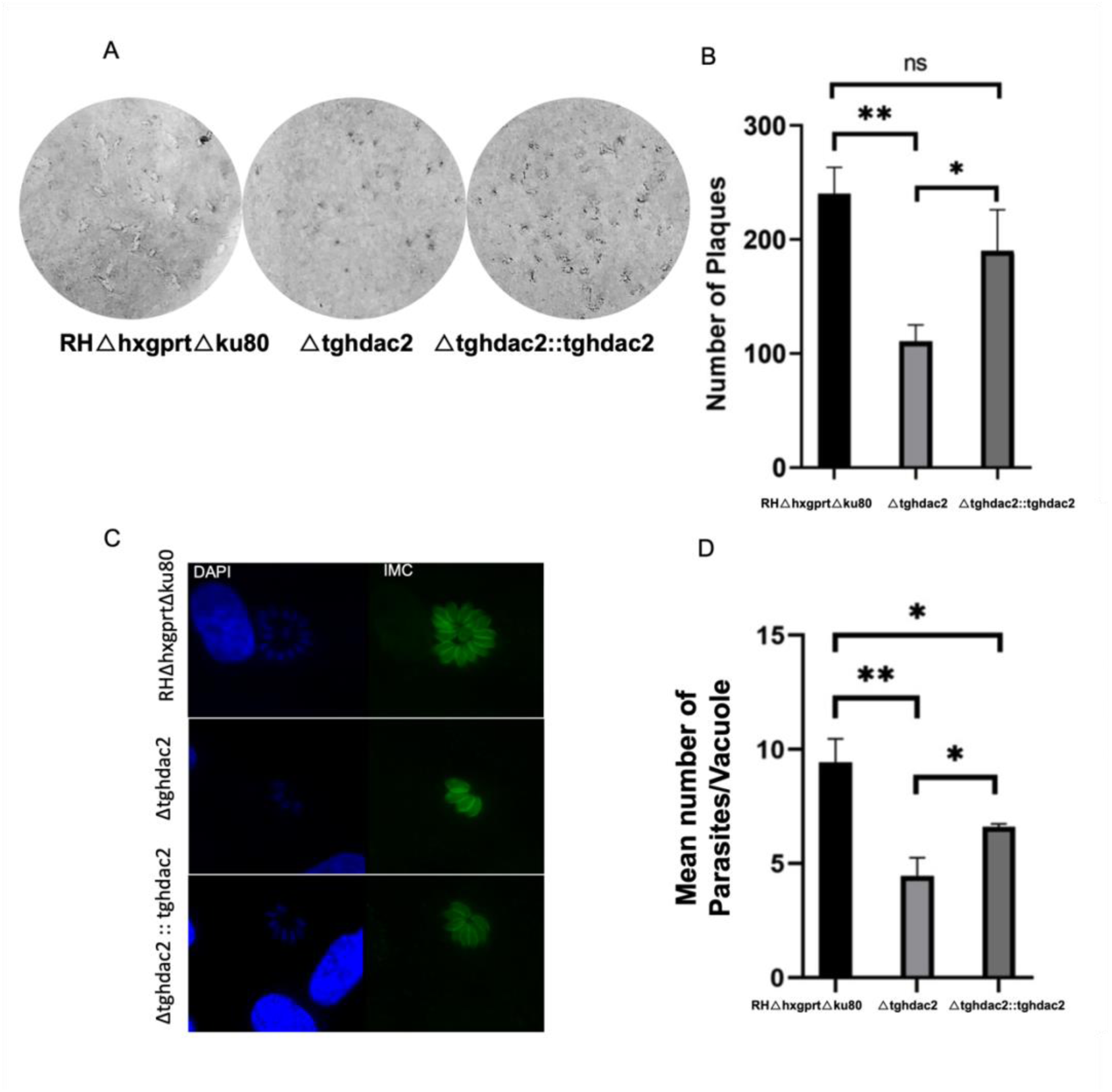
Knockout of *tghdac2* impairs parasite proliferation. (A) Representative image of plaque assay showing the formation of plaques after 500 parasites were added to an HFF monolayer and incubated for five days. Plaques were then counted to assess infection efficiency. (B) Quantification of plaque assay results. (C) Parasites were allowed to infect host cells for one hour, followed by washing with PBS. After 24 hours, the cells were fixed and stained with anti-IMC to visualize the parasites and DAPI to visualize both the nuclei of the parasites and host cells. The number of parasites per vacuole was then counted. (D) Quantification of parasites per vacuole after 24 hours of replication. Data represent the mean of three independent experiments. Statistical analysis was performed using a 2-way ANOVA followed by Tukey’s multiple comparisons test. *P < 0.05, **P < 0.01, ns = not significant.

**Figure 7.**
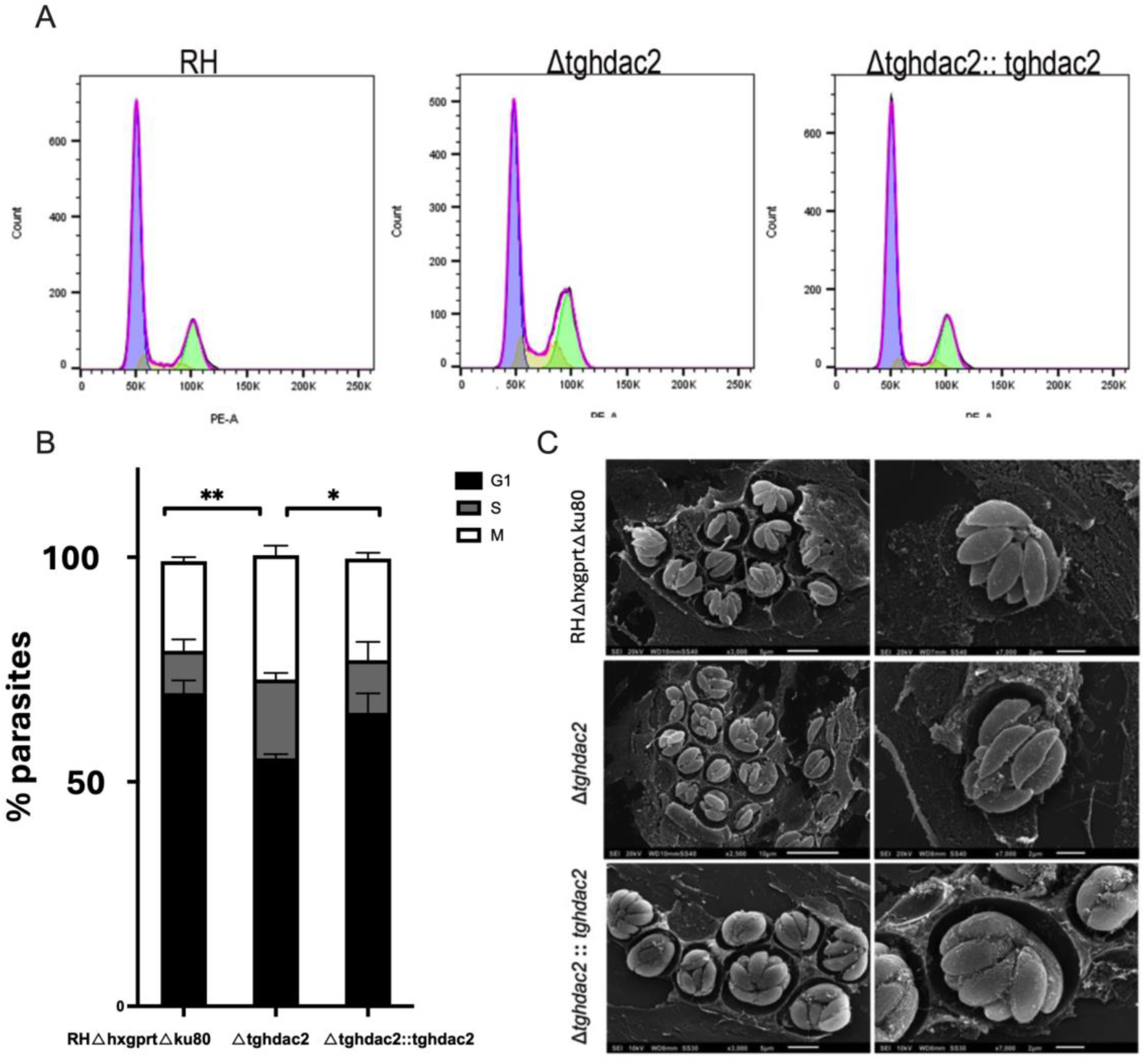
*Tghdac2* knockout parasites exhibit a delay in the S/M phase. (A) Cell cycle analysis was performed 48 hours post-infection. Parasites were purified from host cells, fixed with ethanol, stained with propidium iodide, and analyzed by flow cytometry. The G1 phase is represented in purple, the S phase in yellow, and the G2/M phase in green. (B) Quantification of the cell cycle stages from flow cytometry data. (C) Scanning electron microscopy images 24 hours post-infection show that *tghdac2* knockout parasites are disorganized within the parasitophorous vacuole, with the residual body notably absent. Data are presented as the mean from three independent experiments. Statistical analysis was performed using a 2-way ANOVA followed by Tukey’s multiple comparison test. *P<0.05, **P<0.01. ns = not significant.

Cell cycle disorder can affect the organization of *Toxoplasma* within the parasitophorous vacuole. Scanning electron microscopy confirmed the impact of the *tghdac2* knockout during replication, leading to disorganization of *Δtghdac2* parasites inside the vacuole (Fig. 7C). This phenotype has been previously described as a replication-related defect (38). The knockout effect was partially reverted by complementation of the *tghdac2* gene; however, it did not fully recover the phenotype, likely due to gene not being under the same promoter.

### 3.4 Knockout of *tghdac2* leads to a deregulation of membrane proteins

Knockout parasites showed defects in replication; however, the underlying cause of these defects required further investigation. To explore potential mechanisms, we performed RNA sequencing on the knockout parasites, as well as complemented and parental strains. The data revealed that, on average, 80% of input reads were correctly assigned to genes (Supp. table 3), and 8,638 genes with nonnull counts were identified. We detected 142 differentially expressed genes between the *Δtghdac2* strain and the wild type (Fig. 8A, Supp. Table 4).

**Figure 8.**
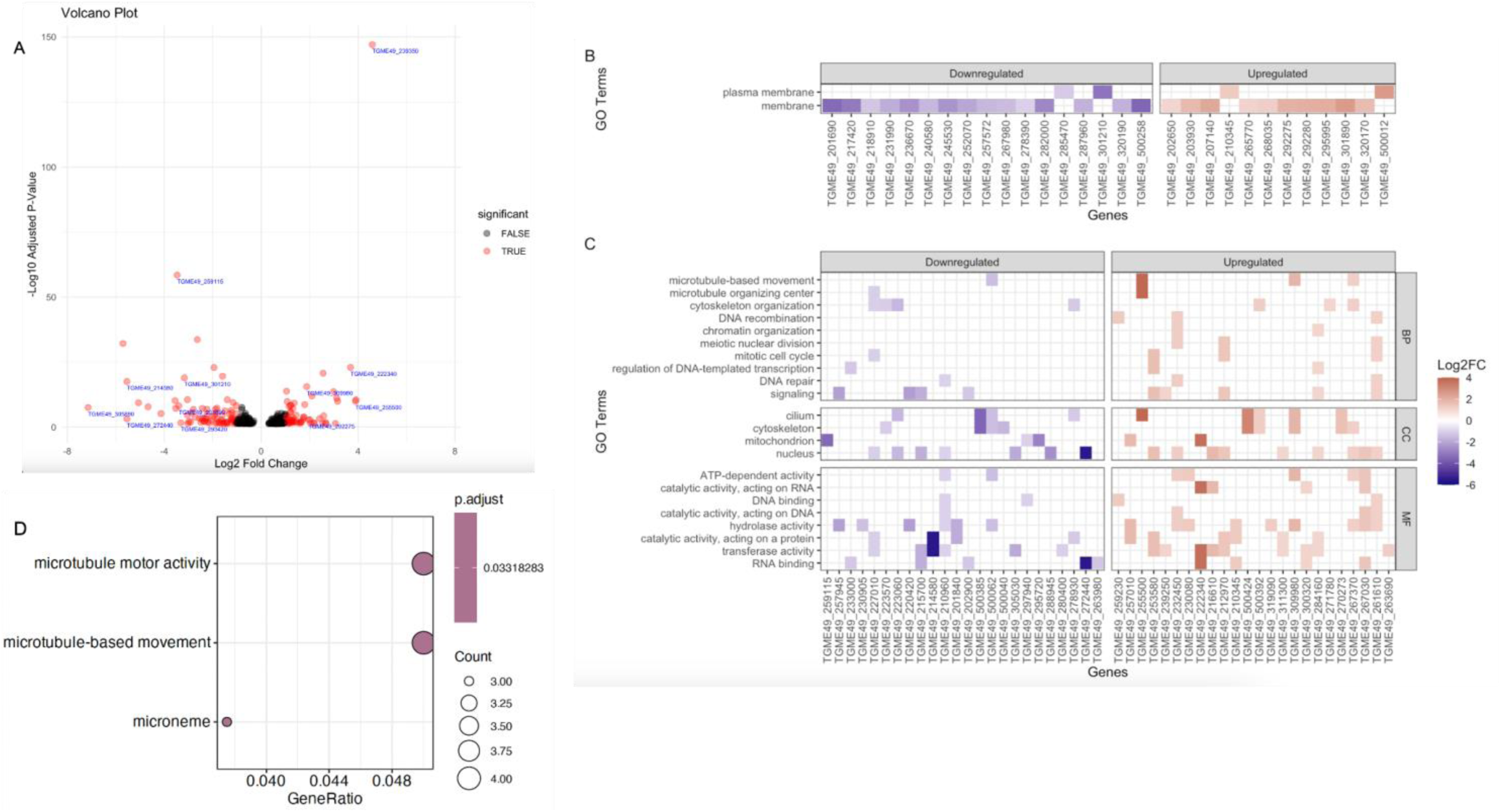
Transcriptome analysis of *T. gondii* strains RH*ΔhxgprtΔku80* and RH*Δtghdac2*. (A) Volcano plot depicting differentially expressed genes (DEGs) between wild-type and knockout parasites. DEGs are highlighted in red, and non-significant genes are shown in black, as determined by the p-value adjusted criterion (< 0.05). For ease of interpretation, the labels of selected genes from Table 2 were added to the plot in blue. Gene ontology (GO) characterization of DEGs depicting (B) membrane-related genes and (C) cytoskeleton/microtubule and DNA/chromatin/signaling-related genes. (D) GO enrichment analysis of the complete set of DEGs.

**Table 2.** Top differentially expressed genes (DEGs). The top DEGs identified in the transcriptome analysis include proteins involved in binding, movement, cell cycle, and membrane transport.

| Gene | FC | Name | Fitness |
| --- | --- | --- | --- |
| TGME49_239350 | 4.59 | RAP domain-containing protein (RNA-binding domain) | -5.13 |
| TGME49_255500 | 3.89 | TPR-like tetratricopeptide-like helical domain superfamily | 0.57 |
| TGME49_222340 | 3.69 | NOL1/NOP2/Sun family protein | -1.28 |
| TGME49_240340 | 2.68 | *Toxoplasma gondii* family E protein | 0.15 |
| TGME49_269310 | 2.64 | Zinc finger (CCCH type) motif-containing protein | -1.07 |
| TGME49_292275 | 1.95 | SAG-related sequence SRS36E | unknown |
| TGME49_309980 | 1.88 | Dynein heavy chain family protein | 0.05 |
| TGME49_305890 | -7.15 | PT repeat-containing protein, putative | -1.01 |
| TGME49_272440 | -5.55 | RNA recognition motif-containing protein | 0.39 |
| TGME49_214580 | -5.54 | Tetratricopeptide repeat-containing protein / Predicted RNA methylase | -0.04 |
| TGME49_259115 | -3.5 | ABC1 family protein | -0.39 |
| TGME49_201690 | -3.40 | BT1 family protein | -1.28 |
| TGME49_301210 | -3.17 | NAD(P) transhydrogenase subunit beta, putative | -1.91 |

Around 70 genes of the DEGs were hypothetical proteins with no identified conserved domains. The top 10 DEGs (excluding hypothetical proteins) included proteins involved in cell cycle progression, binding, and cytoskeleton composition (Table 2). Consistent with this, gene ontology (GO) characterization of all DEGs revealed three main functional categories (Fig. 8B and C, Supp. Fig. 3): membrane, microtubule/cytoskeleton, and broader categories related to the nucleus/DNA/chromatin/cell cycle. Functional enrichment analysis detected three significant microtubule terms (Fig. 8D). These results support our aforementioned findings, suggesting that TgHDAC2 is involved in cell cycle regulation and IMC biogenesis.

Furthermore, the data suggest that the peptide insertion domain plays a role as a cyclin G kinase or microtubule cross-linking factor, both of which are crucial for cell cycle progression and overall microtubule organization within the cell. Additionally, the predominant GO term related to cellular components for both up and downregulated genes was membrane, reinforcing the positioning of TgHDAC2 in the IMC membrane (Fig. 8B).

Finally, to gather further insight into the hypothetical proteins, we searched for conserved domains and families using InterPro Scan (IPS) through the Omics Box software (Supp. Fig. 3). The most enriched families identified included the P-loop containing nucleoside triphosphate hydrolase family (nucleoside binding proteins), the SRS domain superfamily (prevalent on the parasite’s surface), and the S-adenosyl-L-methionine-dependent methyltransferase superfamily. Additionally, dynein domains were detected, including the heavy chain, C-terminal, N-terminal, AAA lid domain, and linker subdomain 3. These findings emphasize a global deregulation of cytoskeleton-interacting genes in the transcriptomic profile of the mutant strain.

## 4 Discussion

The structural conformation of proteins is intrinsically linked to their functionality. According to http://ToxoDB.org and the Conserved Domains Database (CDD, NCBI), although TgHDAC2 contains a characteristic class I HDAC domain, it also contains two peptide insertions within the classical class I HDAC domain. Previous analyses conducted by our group have demonstrated that these insertions are unique to a few members of the phylum Apicomplexa (*T. gondii* and *Hammondia hammondi*), although their specific function or structure remains unknown (18). Based on our current data, this region likely forms a distinct domain (Fig. 4). In this study, we demonstrate that TgHDAC2 is localized to the *Toxoplasma* membranes, where it colocalizes with IMC. This suggests that, in addition to its classification as an HDAC, this protein has different targets.

To further investigate the role of these peptide insertions, a CD analysis was performed to assess their impact on protein stability. Full-length TgHDAC2 failed to renature during thermal denaturation, while TgHDAC2^Δ74-276^ was able to successfully renature. This suggests that the peptide insertions serve a functional role rather than a structural one in the protein.

Supporting this idea, our XL-MS model further suggests that the peptide insertions form a distinct domain, indicating a potential additional functional role. To explore this function, we employed the ProFunc server, which predicts protein function by comparing the 3D structure to known proteins and functional motifs. The ProFunc data suggest a correlation between the peptide insertions and proteins such as cyclin-G-associated kinase (GAK), microtubule crosslinking factor 1 (MACF1), serine/arginine repetitive matrix protein 2, and myb-related transcription factor.

MACF1 is a cytoskeleton component involved in several cellular functions in mammals, including cell proliferation, migration, and signal transduction (39). GAK is essential for clathrin-mediated membrane trafficking and plays an important role in cell cycle progression (40) (41)(42). During the cell cycle, GAK is involved in centrosome maturation, as well as chromosome and mitotic spindle formation (44). Although *T. gondii* lacks GAK, it possesses tyrosine kinase-like (TKL) proteins, which are considered orthologous of mammalian GAK. TLKs have several functions in essential *T. gondii* processes (43).

Additionally, the peptide insertions share structural similarities with Myb-related transcription factor. In *T. gondii,* a Myb-like transcription factor, BFD1, was identified by Waldman et al. (44) as important for differentiating tachyzoites from bradyzoites in culture. The authors found that BFD1 binds to transcription initiation sites of genes expressed in bradyzoites, and its absence impairs the parasite’s ability to differentiate under alkaline stress. Although these predicted functions are plausible, further studies are needed to fully elucidate the role of this additional domain in TgHDAC2.

Another potential function of the peptide insertions could be regulation. Protein activity is commonly regulated by PTMs, and the peptide insertions contain predicted sites for CK2 and PKC, myristoylation, and glycosylation, all of which could influence its structure or functionality. CK2 phosphorylation sites in the C-terminal region of HDACs from mammals and other eukaryotes are known to regulate protein activity (45). For instance, HsHDAC2 is hyperphosphorylated during mitosis, which correlates with increased activity (46)(47)(48). Our *in silico* analysis identified multiple PTMs, suggesting that this region may serve as a regulatory domain for TgHDAC2 activity. This is supported by experimental evidence from the literature, which has reported phosphorylation at serine residues 85, 150, 602, and 603, as well as threonine 179, in TgHDAC2 (49)(50).

In line with our findings, the RNA sequencing and microscopy data suggest that TgHDAC2 interacts with membrane and cytoskeleton proteins within the IMC during its biogenesis and the cell cycle. In addition to serving as a scaffold during endodyogeny, the IMC plays a pivotal role in anchoring essential invasion-related proteins (51) (52) (53).

Fluorescence microscopy images revealed that, prior to the visibility of the nucleus in typical mitotic progression, TgHDAC2 is diffusely distributed throughout the cytoplasm. Accumulation of the protein was observed near the nucleus, potentially indicating the formation of daughter cell scaffolds. This process involves the duplication of centrioles and the positioning of IMC15 and MORN1 near them. Notably, the absence of MORN1 disrupts the formation of the basal complex, impairing the efficient segregation of the apicoplast and daughter cells, leading to a failure in cytokinesis (54).

Although cytokinesis was not impaired in the *Δtghdac2* strain, parasites showed a delay in the cell cycle with accumulation in the S phase. Additionally, the residual body was absent, and the parasites exhibited a disorganized appearance within the vacuole. MORN-domain-containing proteins were found to be downregulated in the transcriptomic analysis (Supplemental Table 4). The disorganization of the parasitophorous vacuole strongly suggests disruptions in the cell cycle. Vacuole disorganization associated with cell cycle disruptions was also observed in *prmt1* knockout parasites (38).

In the M phase, the cytoskeleton expands, with the highest concentration of IMCs in the daughter cells, except for IMCs 5, 8, 9, 13, and centrin 2, which migrate to the basal complex and begin contracting for complete cell division, culminating in cytokinesis (55)(56)(57)(58)(59). During these final stages of the cell cycle, TgHDAC2 extensively surrounds the nucleus, from the initiation of budding to the completion of duplication, colocalizing with daughter IMC proteins. However, it is absent after cytokinesis in the new G1 phase (Fig. 5A). It is plausible to speculate that TgHDAC2 has a role in recruiting proteins to the IMC during its biogenesis.

In addition to IMC proteins, microtubules are essential for the segregation of chromosomes, organelles, and centrioles. Studies have shown that drugs inhibiting microtubules disrupt the formation of the internal membrane complex and the conoid of daughter cells (60). Furthermore, cytoskeletal components like tubulins are highly acetylated. Acetylation at lysine 40 of α-tubulin is enriched during daughter cell formation and is critical for stabilizing microtubules (61). TgATAT has been identified as the acetyltransferase responsible for K40 acetylation in tubulins during the cell cycle (61).

In humans, HDAC6 deacetylates α-tubulin, and tubulin acetylation regulates tubulin binding and kinesin-1 motility (62). Beyond tubulin, IMC proteins have predicted acetylation sites, with 26% of acetylated proteins in *T.gondii* identified as cytoskeletal components (63). This suggests that TgHDAC2 might play a role in deacetylating these proteins throughout the cell cycle. However, additional experimental evidence is required to confirm this hypothesis.

Furthermore, the gene expression profile analysis of the knockout parasites supports a role in membrane proteins and cytoskeletal components. Most differentially expressed proteins are membrane-associated, with many also identified as cytoskeletal proteins.

In addition, one of the top hits in downregulated genes is TGME49_305890, a PT-repeat protein containing a DNA translocase FtsK domain. In bacteria, this domain represents the fastest DNA translocation motor identified to date. It is essential for bacterial cell division, acting in cytokinesis, chromosome segregation, and daughter cell separation (64) (65) (66) (67). While there is no information on acetylation in FtsK proteins, the downregulation of this protein in *Δtghdac2* parasites may explain the observed delay in the cell cycle.

Another noteworthy finding is the presence of genes containing a tetratricopeptide-like (TPR-like) domain, found among the top hits of DEGs (TGME49_255500 and TGME49_214580). This domain is versatile, contributing to various functions, including the formation of multi-protein complexes, gene expression regulation, and cell cycle progression. Some members of this domain family are involved in mitosis, serving as a critical component for cell division and cytokinesis(68).

In summary, our findings revealed that TgHDAC2 plays a pivotal role in the cell cycle of *T. gondii*, potentially facilitating IMC biogenesis through tubulin deacetylation or other essential proteins required for cell division. This study broadens our understanding of the parasite’s biology, highlighting that the function of *T.gondii* HDACs extends beyond regulating gene expression. Additionally, we resolved a previously uncharacterized structure, suggesting that TgHDAC2 operates in a unique and specialized manner in *T. gondii*. These insights provide valuable perspectives on key biological processes critical for the parasite’s survival and may guide the development of therapeutic strategies targeting this unique HDAC function.

## Acknowledgements

We thank the Program for Technological Development in Tools for Health-RPT FIOCRUZ for Technological Platforms of Flow Cytometry, Microscopy, Mass Spectrometry, and Protein Purification. We also would like to thank Gary Ward and Peter Bradley for kindly providing antibodies and Beatriz Borges for scanning electron microscopy pictures. This research used facilities of the Brazilian Biorenewables National Laboratory (LNBR), part of the Brazilian Centre for Research in Energy and Materials (CNPEM), a private non-profit organization under the supervision of the Brazilian Ministry for Science, Technology, and Innovations (MCTI). The High-Performance Sequencing (SEQ) open access facility staff is acknowledged for the assistance during the experiments (proposal 20220471).

S.N. was supported by grants from Fiocruz (VPPCB-FIO-007-FIO-18-2-92), CNPq (PROEP-ICC 442339/2019-4), Fundação Araucária-CNPq (PI 01/2018), CAPES-COFECUB (88881.370870/2019-01) and resources from Instituto Carlos Chagas, Fiocruz.

## Supplemental Material

**Supplemental Table 1.**
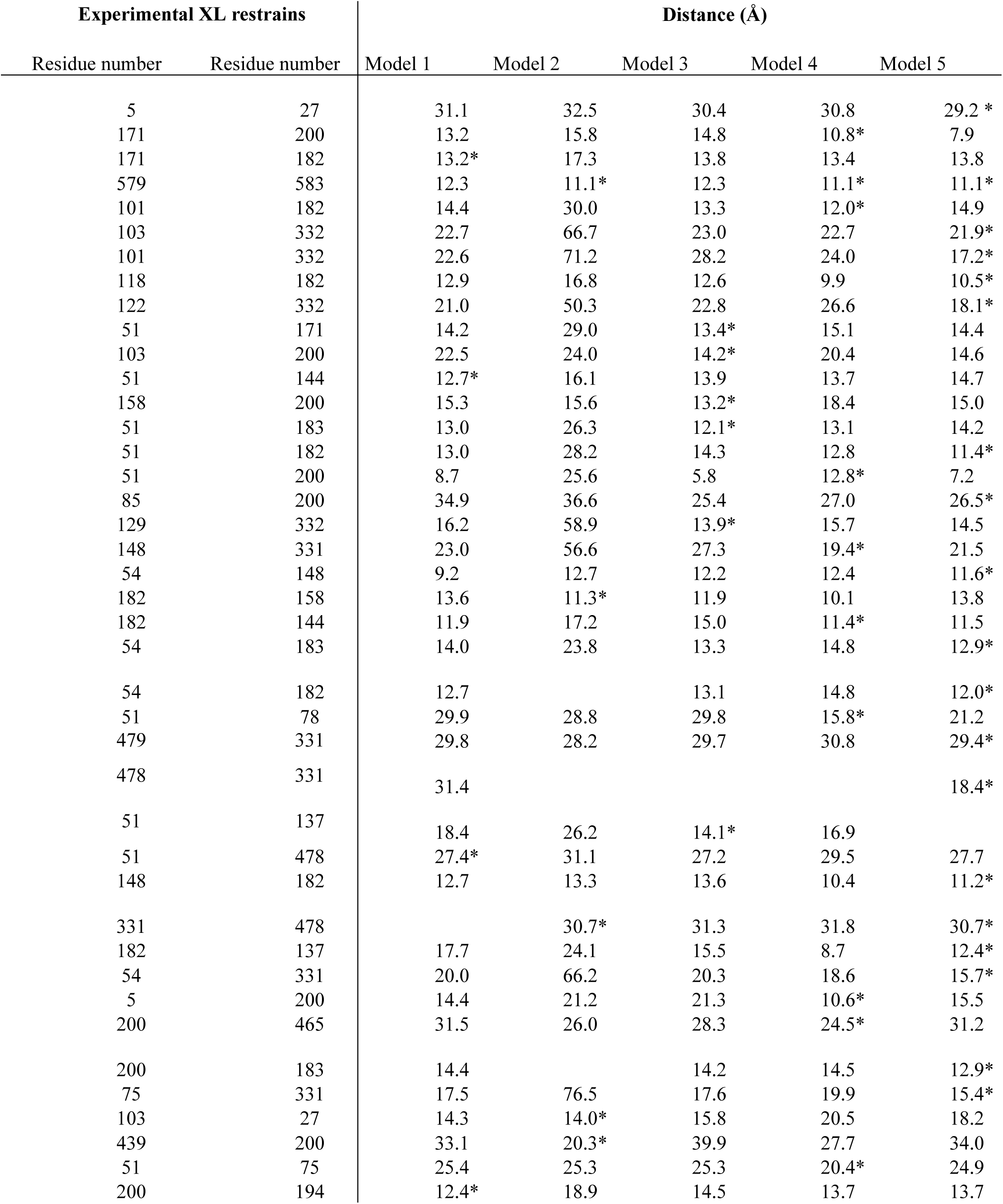
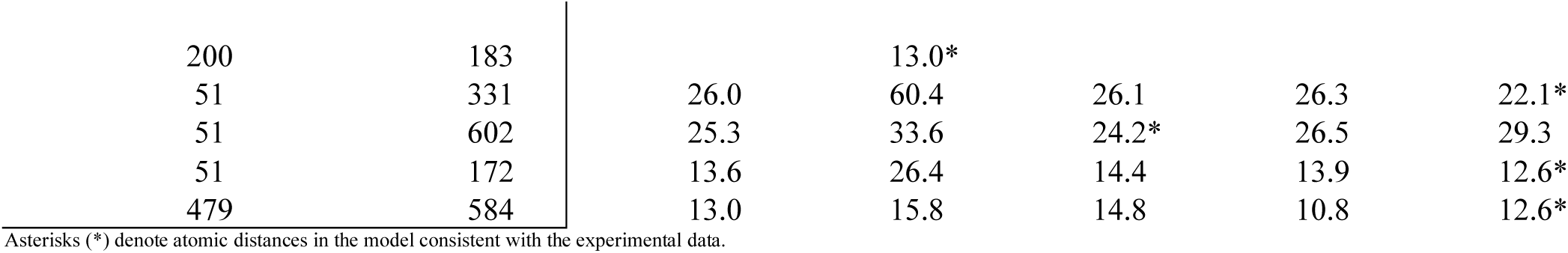
XL restraints obtained by XLMS and respective spatial distances between Ca in three-dimensional models.

| Experimental XL restrains |  | Distance (Å) |  |  |  |  |
| --- | --- | --- | --- | --- | --- | --- |
| Residue number | Residue number | Model 1 | Model 2 | Model 3 | Model 4 | Model 5 |
| 5 | 27 | 31.1 | 32.5 | 30.4 | 30.8 | 29.2 * |
| 171 | 200 | 13.2 | 15.8 | 14.8 | 10.8* | 7.9 |
| 171 | 182 | 13.2* | 17.3 | 13.8 | 13.4 | 13.8 |
| 579 | 583 | 12.3 | 11.1* | 12.3 | 11.1* | 11.1* |
| 101 | 182 | 14.4 | 30.0 | 13.3 | 12.0* | 14.9 |
| 103 | 332 | 22.7 | 66.7 | 23.0 | 22.7 | 21.9* |
| 101 | 332 | 22.6 | 71.2 | 28.2 | 24.0 | 17.2* |
| 118 | 182 | 12.9 | 16.8 | 12.6 | 9.9 | 10.5* |
| 122 | 332 | 21.0 | 50.3 | 22.8 | 26.6 | 18.1* |
| 51 | 171 | 14.2 | 29.0 | 13.4* | 15.1 | 14.4 |
| 103 | 200 | 22.5 | 24.0 | 14.2* | 20.4 | 14.6 |
| 51 | 144 | 12.7* | 16.1 | 13.9 | 13.7 | 14.7 |
| 158 | 200 | 15.3 | 15.6 | 13.2* | 18.4 | 15.0 |
| 51 | 183 | 13.0 | 26.3 | 12.1* | 13.1 | 14.2 |
| 51 | 182 | 13.0 | 28.2 | 14.3 | 12.8 | 11.4* |
| 51 | 200 | 8.7 | 25.6 | 5.8 | 12.8* | 7.2 |
| 85 | 200 | 34.9 | 36.6 | 25.4 | 27.0 | 26.5* |
| 129 | 332 | 16.2 | 58.9 | 13.9* | 15.7 | 14.5 |
| 148 | 331 | 23.0 | 56.6 | 27.3 | 19.4* | 21.5 |
| 54 | 148 | 9.2 | 12.7 | 12.2 | 12.4 | 11.6* |
| 182 | 158 | 13.6 | 11.3* | 11.9 | 10.1 | 13.8 |
| 182 | 144 | 11.9 | 17.2 | 15.0 | 11.4* | 11.5 |
| 54 | 183 | 14.0 | 23.8 | 13.3 | 14.8 | 12.9* |
| 54 | 182 | 12.7 |  | 13.1 | 14.8 | 12.0* |
| 51 | 78 | 29.9 | 28.8 | 29.8 | 15.8* | 21.2 |
| 479 | 331 | 29.8 | 28.2 | 29.7 | 30.8 | 29.4* |
| 478 | 331 | 31.4 |  |  |  | 18.4* |
| 51 | 137 | 18.4 | 26.2 | 14.1* | 16.9 |  |
| 51 | 478 | 27.4* | 31.1 | 27.2 | 29.5 | 27.7 |
| 148 | 182 | 12.7 | 13.3 | 13.6 | 10.4 | 11.2* |
| 331 | 478 |  | 30.7* | 31.3 | 31.8 | 30.7* |
| 182 | 137 | 17.7 | 24.1 | 15.5 | 8.7 | 12.4* |
| 54 | 331 | 20.0 | 66.2 | 20.3 | 18.6 | 15.7* |
| 5 | 200 | 14.4 | 21.2 | 21.3 | 10.6* | 15.5 |
| 200 | 465 | 31.5 | 26.0 | 28.3 | 24.5* | 31.2 |
| 200 | 183 | 14.4 |  | 14.2 | 14.5 | 12.9* |
| 75 | 331 | 17.5 | 76.5 | 17.6 | 19.9 | 15.4* |
| 103 | 27 | 14.3 | 14.0* | 15.8 | 20.5 | 18.2 |
| 439 | 200 | 33.1 | 20.3* | 39.9 | 27.7 | 34.0 |
| 51 | 75 | 25.4 | 25.3 | 25.3 | 20.4* | 24.9 |
| 200 | 194 | 12.4* | 18.9 | 14.5 | 13.7 | 13.7 |
| 200 | 183 |  | 13.0* |  |  |  |
| 51 | 331 | 26.0 | 60.4 | 26.1 | 26.3 | 22.1* |
| 51 | 602 | 25.3 | 33.6 | 24.2* | 26.5 | 29.3 |
| 51 | 172 | 13.6 | 26.4 | 14.4 | 13.9 | 12.6* |
| 479 | 584 | 13.0 | 15.8 | 14.8 | 10.8 | 12.6* |

Asterisks (\*) denote atomic distances in the model consistent with the experimental data.

**Supplemental Table 2.**
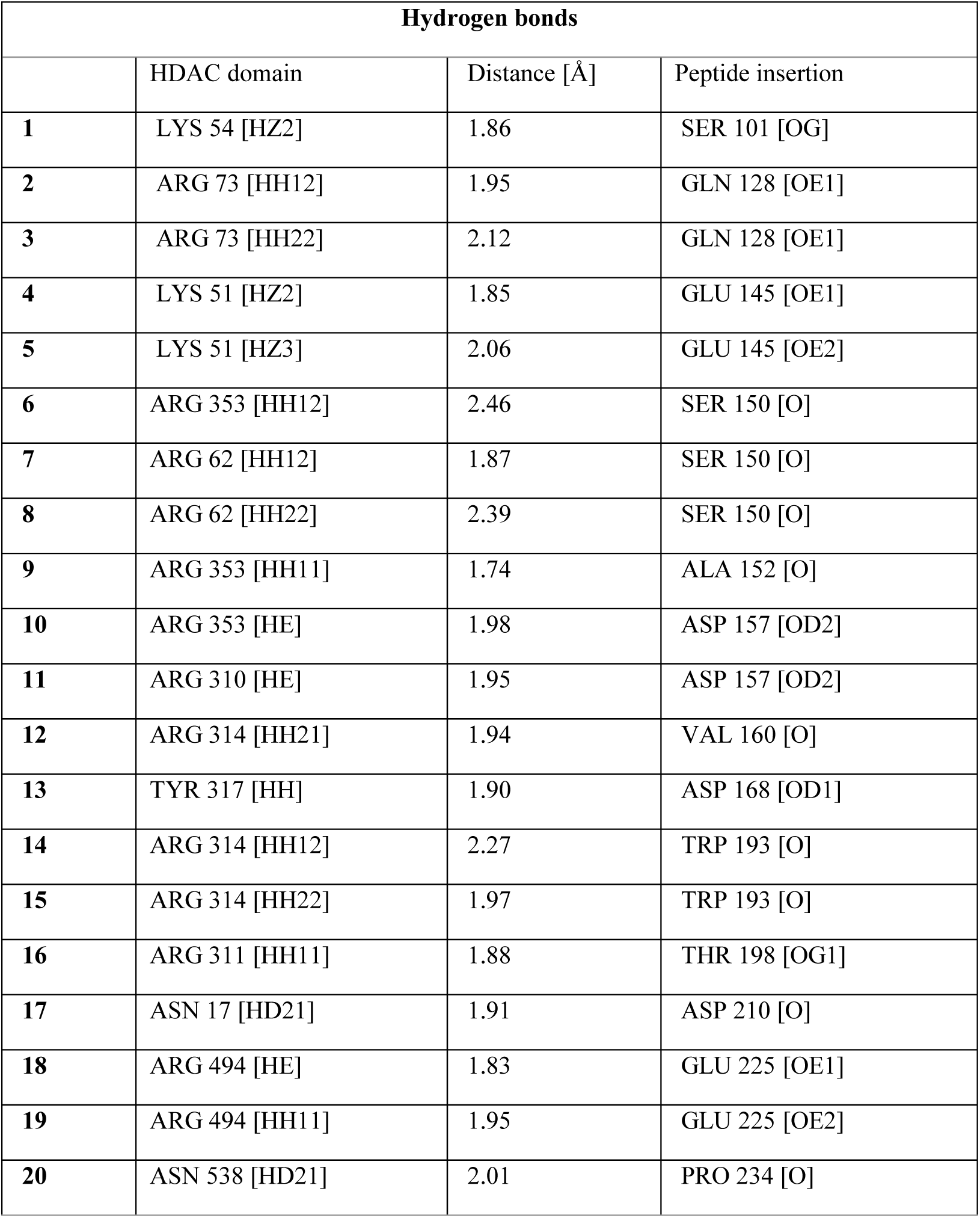
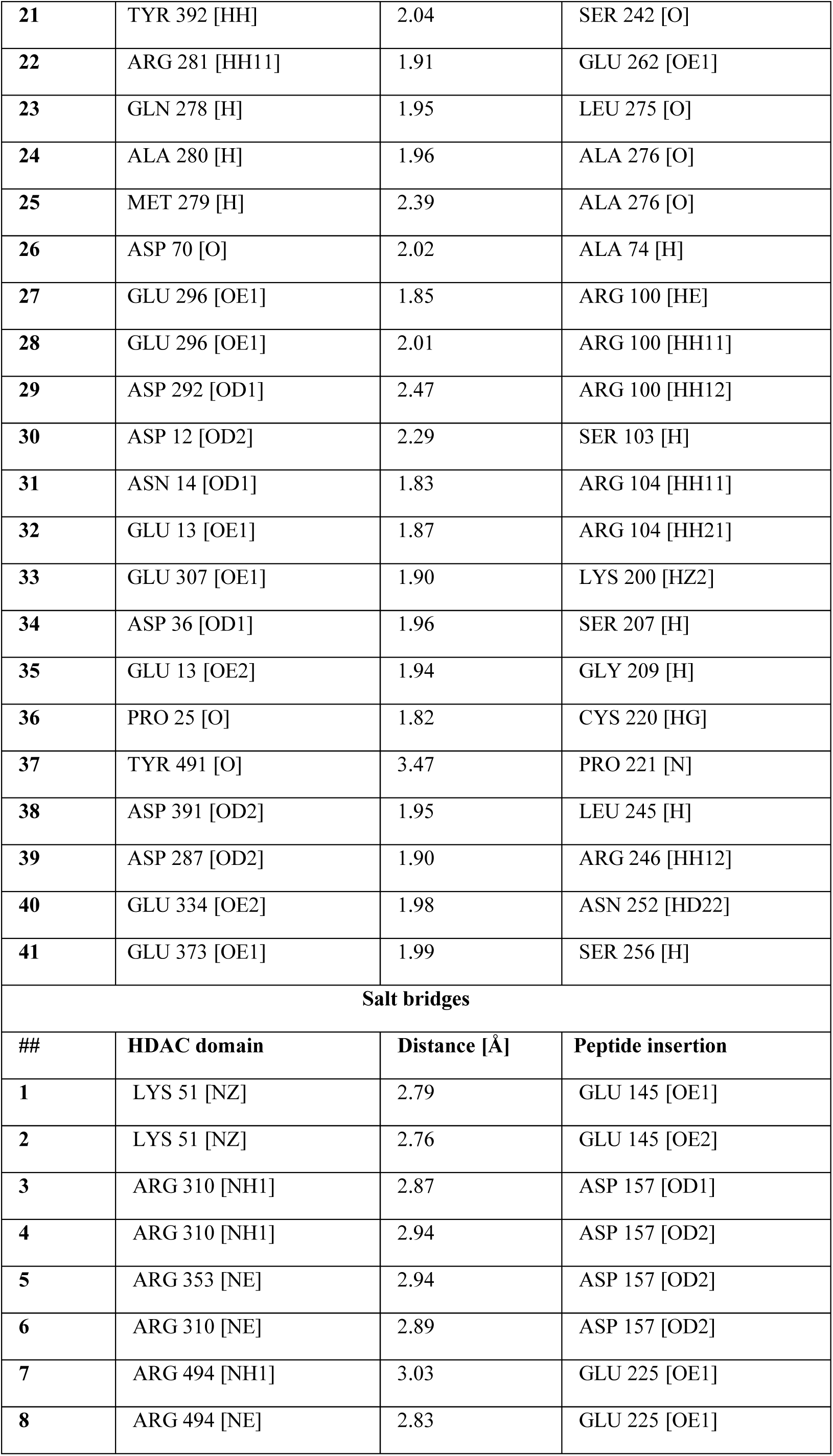
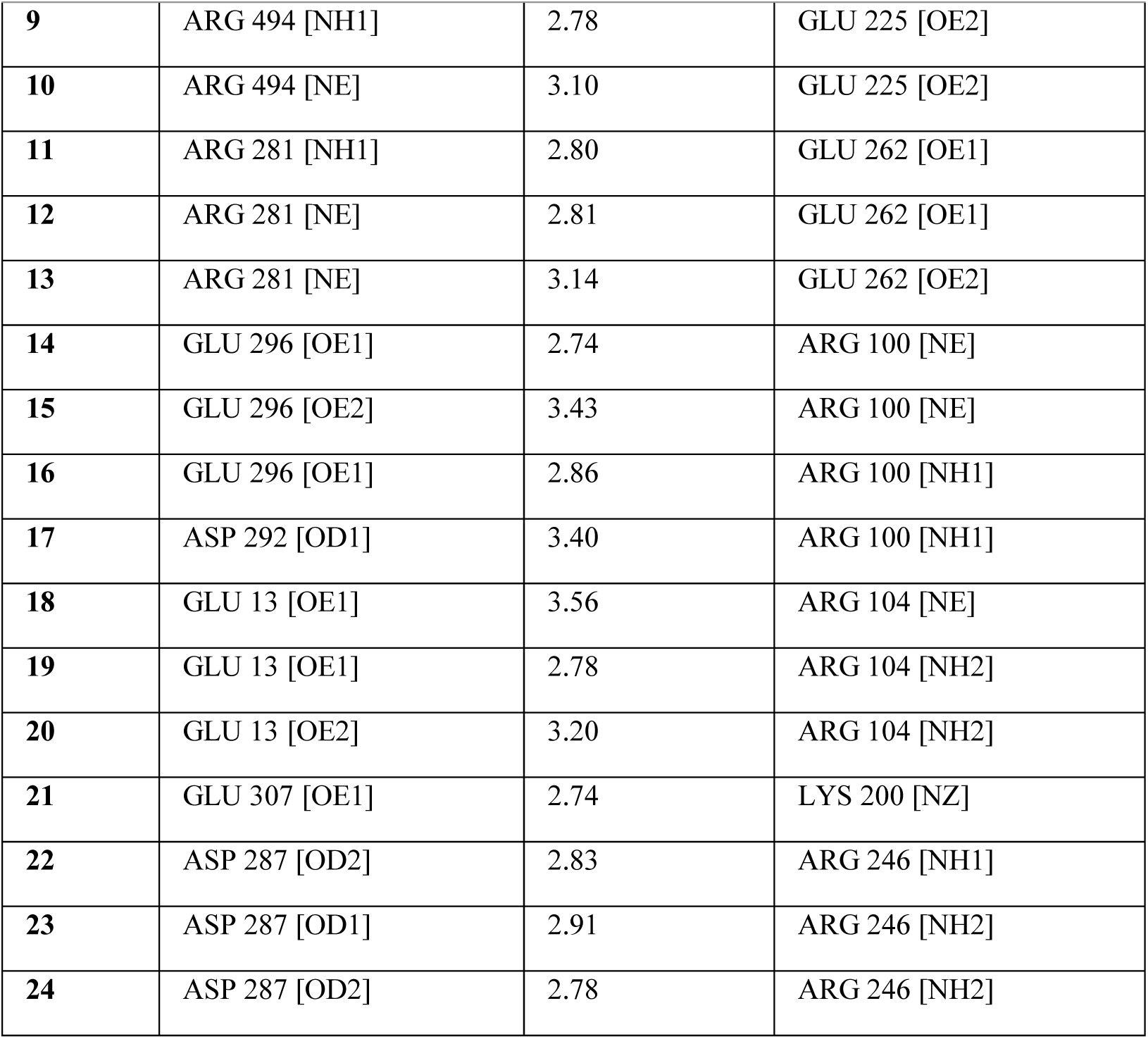
Interaction observed between the HDAC domain and the peptide insertion.

**Supplemental Table 3.** https://docs.google.com/spreadsheets/d/1IlP-PS-7PJOKt144lKyoKQEc3_yPzE34mY8XdEzyCkE/edit?usp=sharing

**Supplemental Table 4.** https://docs.google.com/spreadsheets/d/1qRpmVkbYJHht5DRDomnAW_SWEADzqyV/edit?usp=sharing&ouid=116428869499929794964&rtpof=true&sd=true

**Supplemental Table 5.** Primers. https://docs.google.com/spreadsheets/d/1g0kl8ahDIoPudpEfv8ISbrr5ezI2g3IE/edit?usp=sharing&ouid=116428869499929794964&rtpof=true&sd=true

**Supplemental Figure 1.**
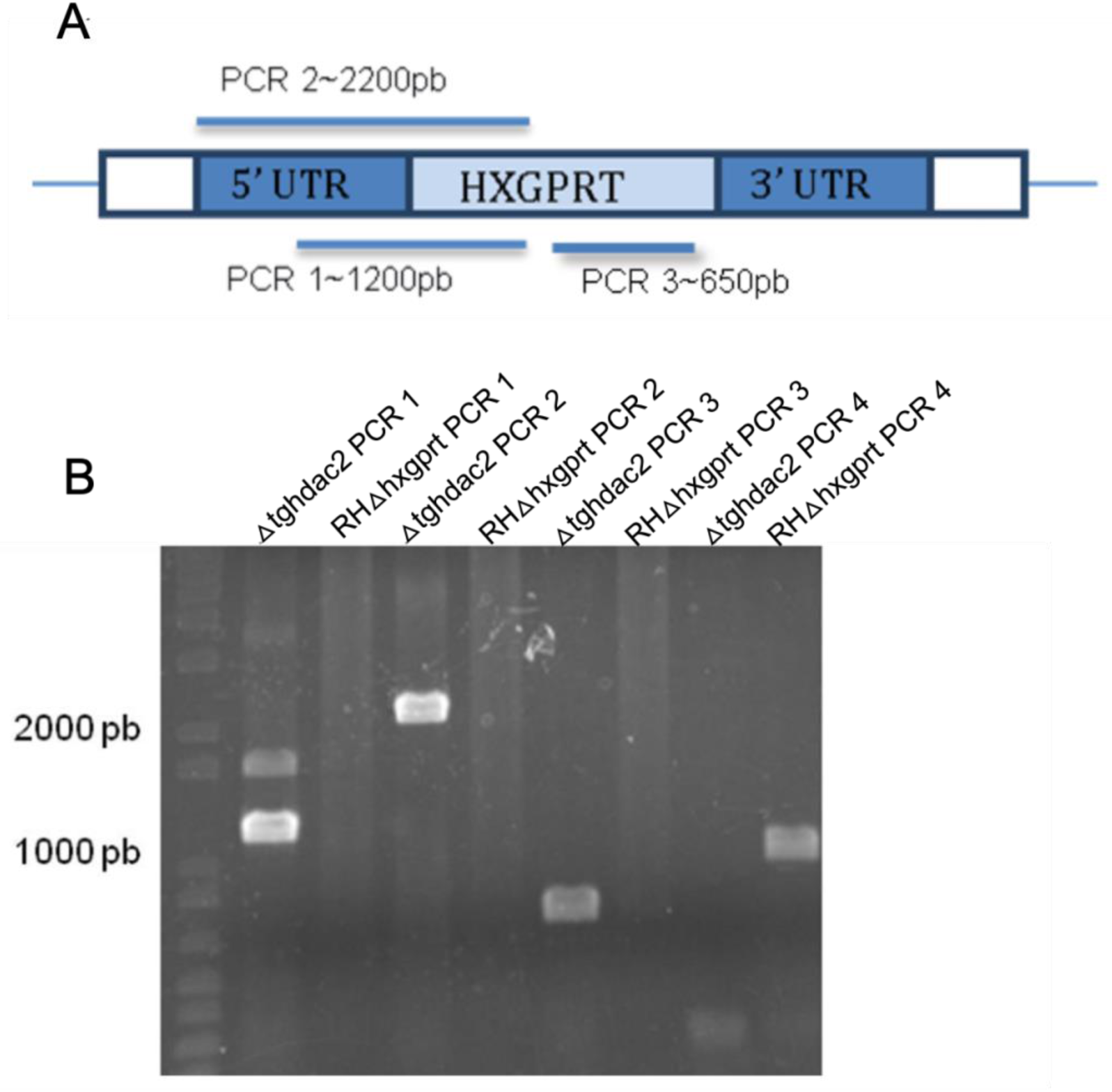
Confirmation of *tghdac2* gene knockout by PCR. (A) Schematic representation of the primers used for PCR amplification at the *tghdac2* locus. The *tghdac2* gene was replaced by the *hxgprt* gene in the knockout clone. (B) Agarose gel electrophoresis of the PCR products, demonstrating successful amplification consistent with the *tghdac2* knockout. Four PCR reactions were performed, including PCR 4, which targets a 1,000 bp region within the *tghdac2* gene. DNA from the knockout clone and the parental *Toxoplasma gondii* strain RHΔhxgprt Δku80 were used as templates. 1 kb Plus DNA Ladder (Thermo Fisher Scientific, Waltham, Massachusetts, USA) was used as a molecular size marker.

**Supplemental Figure 2.**
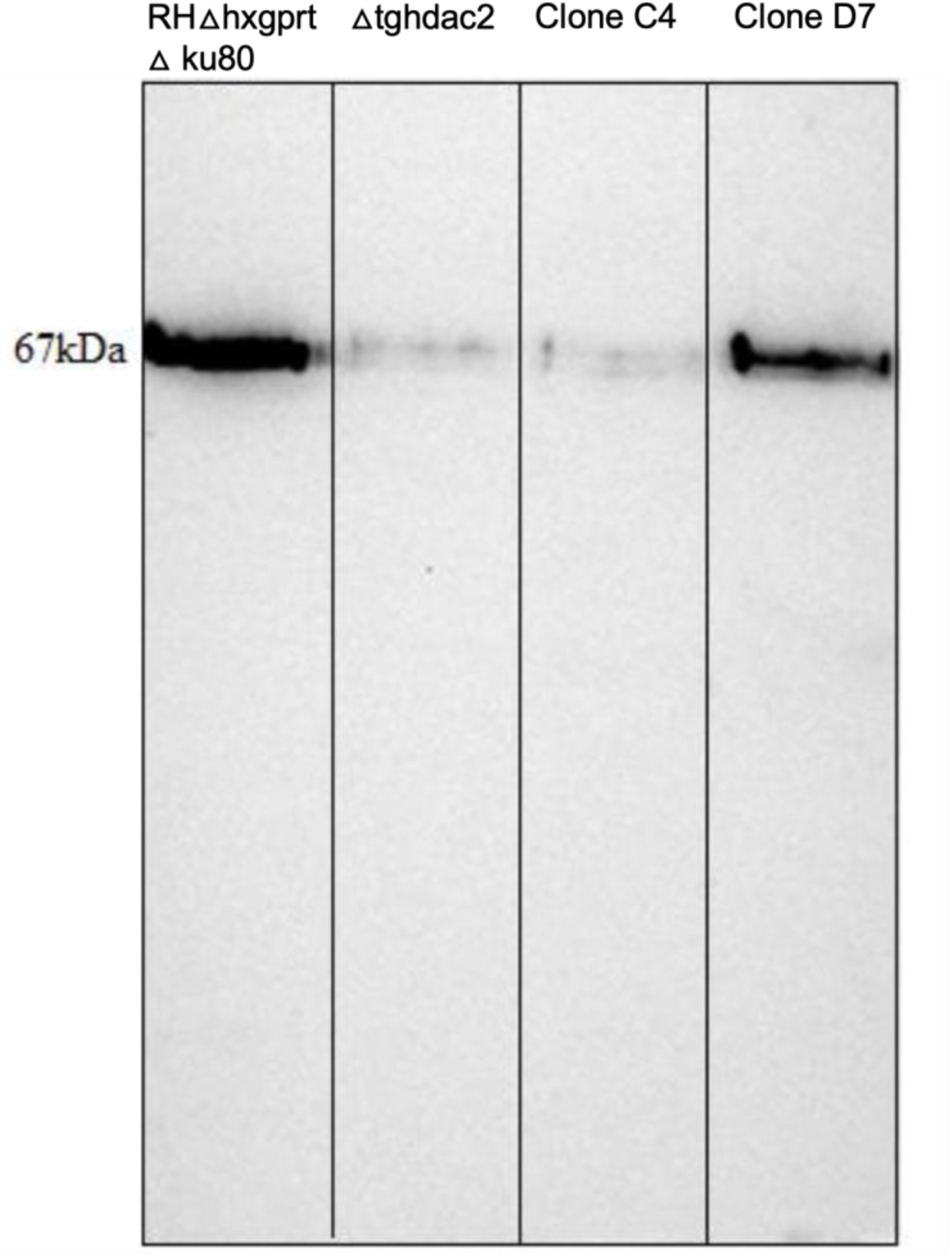
Western blot analysis of TgHDAC2 expression in complemented clones. Western blot analysis was performed using an anti-TgHDAC2 antibody to assess TgHDAC2 expression in complemented clones C4 and D7. The parental *Toxoplasma gondii* strains RHΔhxgprt Δku80 (positive control) and *Δtghdac2* (negative control) were included to confirm the presence or absence of TgHDAC2, respectively. The results demonstrated the expression of this protein in the complemented clone D7.

**Supplemental Figure 3.**
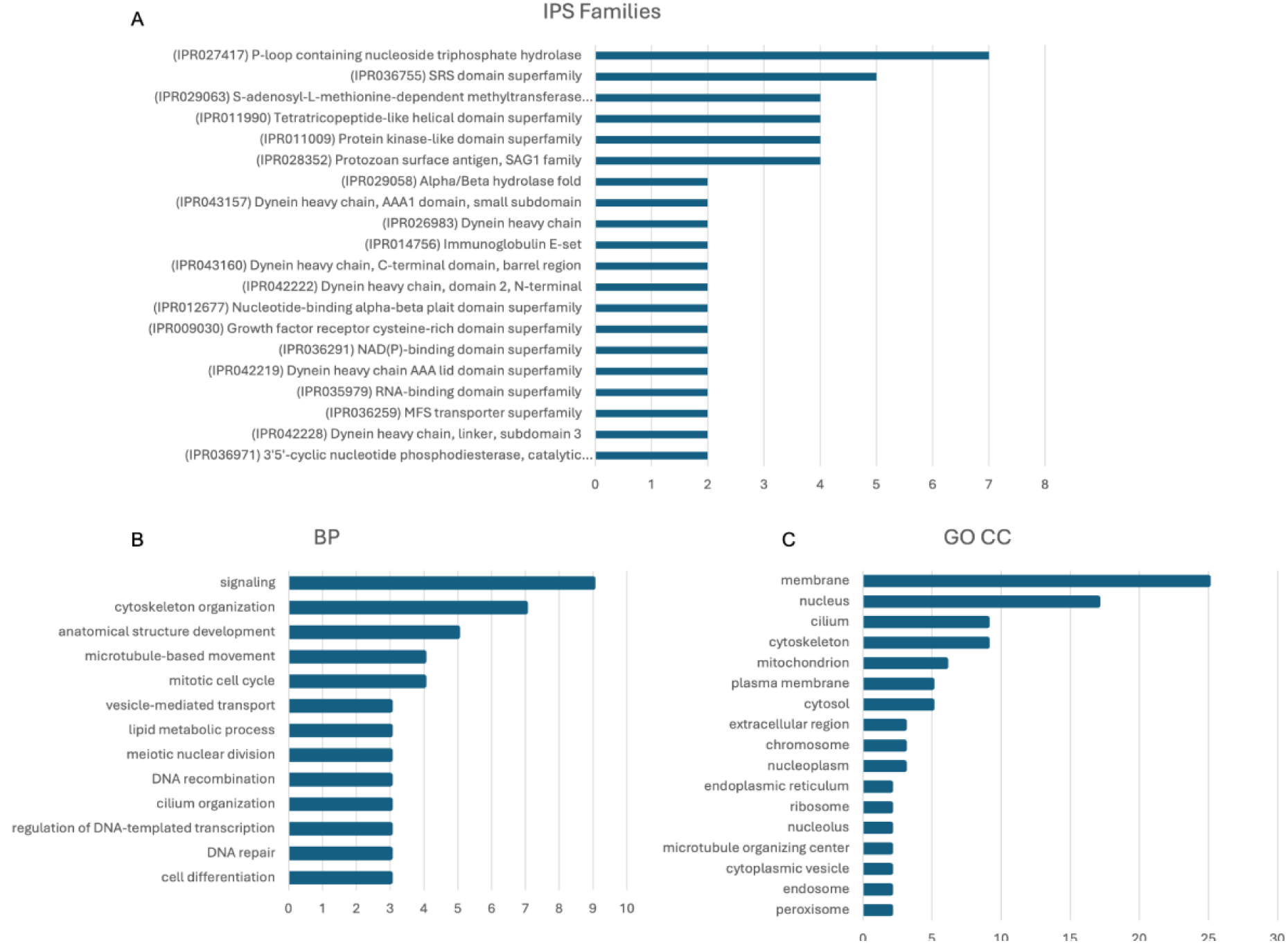
Search for families and gene ontology (GO) terms of differentially expressed genes (DEGs). (A) InterPro Scan (IPS) families identified in DEGs. (B) GO terms related to biological processes. (C) GO terms related to cellular components of DEGs.

